# Facilitating analysis of open neurophysiology data on the DANDI Archive using large language model tools

**DOI:** 10.1101/2025.07.17.663965

**Authors:** Jeremy F. Magland, Ryan Ly, Oliver Rübel, Benjamin Dichter

## Abstract

The DANDI Archive is a key resource for sharing open neurophysiology data, hosting over 400 datasets in the Neurodata Without Borders (NWB) format. While these datasets hold tremendous potential for reanalysis and discovery, many researchers face barriers to reuse, including unfamiliarity with access methods and difficulty identifying relevant content. Here we introduce an AI-powered, agentic chat assistant and a notebook generation pipeline. The chat assistant serves as an interactive tool for exploring DANDI datasets. It leverages large language models (LLMs) and integrates with agentic tools to guide users through data access, visualization, and preliminary analysis. The notebook generator analyzes dataset structure with minimal human input, executing inspection scripts and generating visualizations. It then produces an instructional Python notebook tailored to the dataset. We applied this system to 12 recent datasets. Review by neurophysiology data specialists found the generated notebooks to be generally accurate and well-structured, with most notebooks rated as “very helpful.” This work demonstrates how AI can support FAIR principles by lowering barriers to data reuse and engagement.

## Introduction

Since it was launched in 2020, the DANDI Archive [Rübel 2022; Hawrylycz 2023] has become a central resource for open data in neurophysiology. As of this writing, it holds over 400 neurophysiology “dandisets” (DANDI datasets) in the Neurodata Without Borders (NWB) standard [Rübel 2022; Teeters 2015]. Together, these datasets contain more than 350 TB of neurophysiology data from over 20 species, including intracellular and extracellular electrophysiology, calcium imaging, fiber photometry, and a range of behavioral measures, both naturalistic and task-based. These datasets are the basis for many influential studies in systems neuroscience, and provide insight into many contemporary topics in systems neuroscience, such as memory and navigation; auditory, visual, olfactory, and tactile sensory processing; motor control; and decision making.

Analysis of these open datasets holds enormous potential for scientific discovery [de Vries 2023; Rahimzadeh 2023; Burman 2023; Mendoza-Halliday 2024; Zeisler 2025] and tool development [Schneider 2023; Pachitariu 2024; Stringer 2025], and a core goal of the DANDI project is to promote their use. However, there are barriers to broader engagement with data reuse: many scientists are more familiar with analyzing data that they acquired or working within established collaborations [Tenopir 2011]. Analyzing public datasets requires learning new methods for data access and processing, trust in the quality and completeness of the data, and efficient tools to identify datasets relevant to specific research questions.

While significant effort has gone into developing general tutorials and documentation [DANDI Team, PyNWB, MatNWB] for working with NWB data on DANDI, these resources are often either too broad for investigating a specific dandiset, or too narrowly focused on datasets that may not be relevant to the user’s current goals. The diversity of data on the archive, spanning a wide range of scientific questions, experimental designs, stimuli, and recording techniques, makes it particularly difficult to build tools and guidance that support effective dataset discovery, evaluation, and analysis.

Some tools are already in place to help neuroscientists explore DANDI. Neurosift [Magland 2024] enables interactive exploration of neurophysiology data by streaming directly from the archive and providing modality-specific visualizations. However, it is built in the language of the web browser (TypeScript/JavaScript), which limits accessibility for most neuroscientists. Furthermore, its visualization plugins are generic and not tailored to individual dandisets and specific scientific questions. For more specific guidance, the DANDI team maintains an “example_notebooks” GitHub repository [DANDI Team], where users, often data contributors, share Jupyter notebooks demonstrating how to load, visualize, and sometimes reproduce figures from corresponding papers. Additional resources like the OpenScope DataBook [Ager 2024] offer high-quality tutorials for selected datasets from the Allen Institute OpenScope program. While valuable, these currently cover only a small fraction of the 400+ dandisets available.

Recent advances in large language models (LLMs) offer a promising new approach to lowering the barriers to data reuse by automatically or semi-automatically generating dataset-specific introductory material to help scientists get started with analysis. In addition to coding capabilities [Chen 2021], popular LLMs have the required domain knowledge about neuroscience, in some cases surpassing world experts in the field [Luo 2025]. These models also support natural language interaction, making it possible to build conversational tools that can guide users through the process of dataset evaluation and analysis.

To realize this potential, we developed an AI-powered system consisting of an interactive, agentic chat interface and an automated notebook generator. Users can engage with the system in two complementary ways. They can interact directly with an AI agent through a chat interface, asking questions about a specific dandiset, requesting visualizations, or generating code tailored to that dataset. Alternatively, users can work with a Jupyter notebook produced by the system, which serves as a structured, executable guide for understanding and analyzing the dataset. This notebook is generated by an AI agent that first explores the dataset by inspecting NWB files, analyzing metadata, and producing visualizations. The exploration process supports human intervention at key steps, allowing users to correct errors or guide the analysis if needed. Together, the chat interface and notebooks provide both interactive support and reusable documentation to help researchers engage with public neurophysiology data.

## Results

### Chat interface

The *Dandiset Explorer*, shown in Figure 1, is an interactive chat interface where users can explore a particular dandiset using natural language. The chat agent is equipped with tools to query the DANDI API for dandiset-level metadata and to list the NWB files contained in a given dataset. For individual NWB files, the agent can retrieve a structured summary of their contents, including the types of neurodata present and associated metadata. It can also retrieve Python usage documentation for individual NWB files describing how to programmatically access these objects (see get_nwbfile_info in Methods). The agent executes Python scripts in a sandboxed environment, allowing it to stream data directly from remote NWB files and produce both textual and visual outputs in response to user questions.

**Figure 1.**
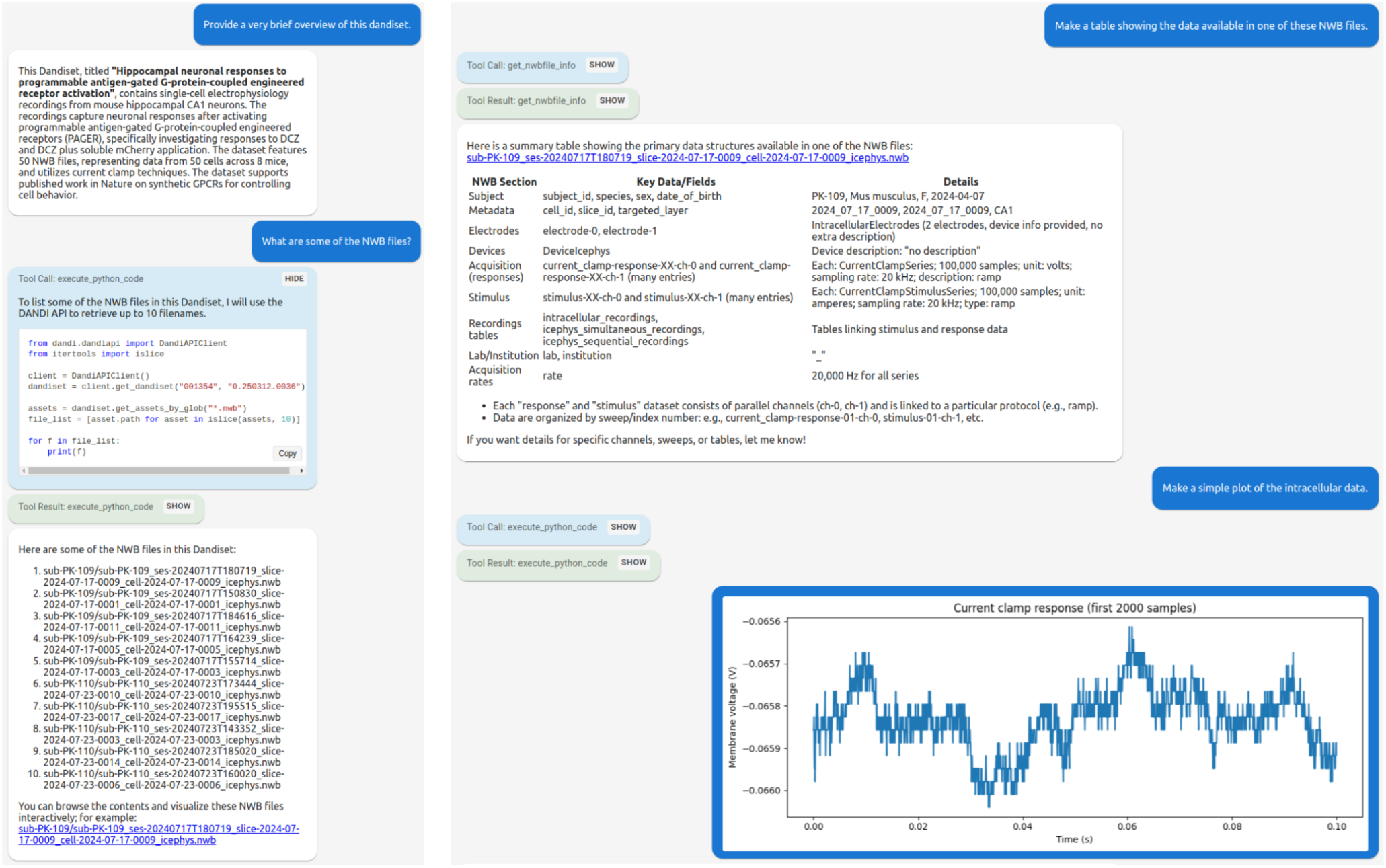
Screenshots of an interactive chat conversation for Dandiset 001354.

### Notebook generation

The notebook generation process (Figure 2) begins with an initial prompt (Appendix A) provided to the chat interface, which explains that the system will conduct a structured exploration to collect the information needed for generating an introductory analysis notebook. This prompt outlines how the exploration should proceed and specifies the criteria for a high-quality notebook. The assistant then proceeds step by step, pausing after each action to wait for user input. If the user replies with “proceed,” the assistant continues autonomously; otherwise, the user can interject at any point with corrections or instructions. This interaction loop enables a flexible balance between automation and human guidance. During exploration, the assistant enumerates NWB files, inspects their structure, and summarizes the types of data present. The agent generates and executes code to create visualizations illustrating key content. If execution raises any exceptions, the resulting error messages are returned to the LLM, which revises the code accordingly. This process can repeat multiple times, providing the agent with multiple attempts to create functioning code. The system is designed to accommodate human oversight in resolving issues such as data inconsistencies or ambiguous metadata. The process continues until the assistant determines that it has gathered sufficient information for notebook generation.

**Figure 2.**
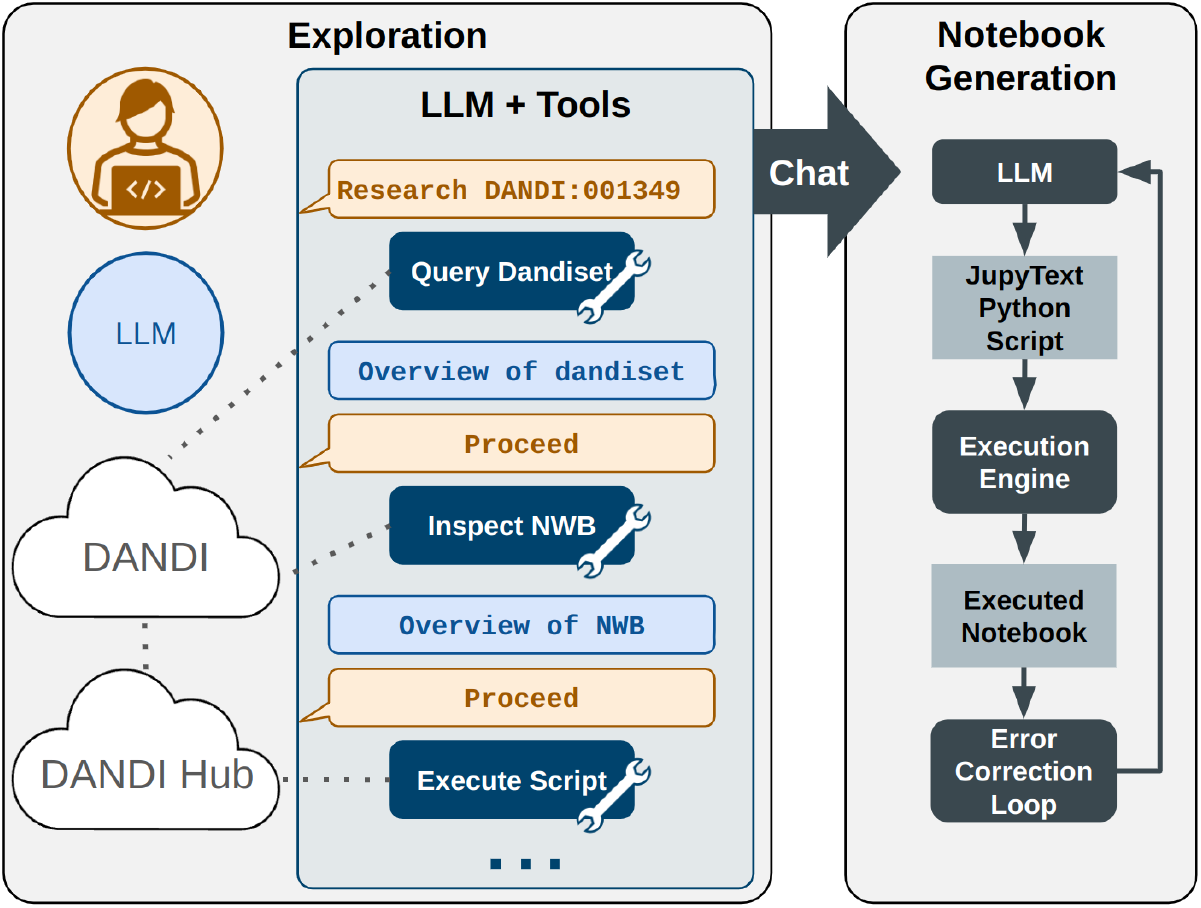
Two-stage process for notebook generation.

Following the exploration phase, a separate LLM agent is tasked with generating a complete Jupyter notebook based on the accumulated interaction history (Appendix A, first prompt). This includes summaries, visual outputs, and code produced during exploration. The LLM is guided by structured instructions (Appendix A, second prompt) to produce the notebook (see Methods).

### Notebooks for twelve published dandisets

To evaluate the system across a variety of real-world datasets, we applied it to 12 recently published dandisets [Huang 2025; Klein 2025; Breathing 2025; Reinagel 2025; Sosa 2025; Ranjan 2025; Gonzalez 2025; Eckert 2025; Galvan 2025; Mehta 2025; Keyes 2025; Berry 2025] spanning a range of species, modalities, and experimental designs. Published dandisets are immutable, in contrast to draft versions, and therefore represent stable targets for notebook generation. Each was processed using the full pipeline: initiating structured exploration via the chat interface, generating a notebook, and executing the result with iterative error correction. Figure 3 shows screenshots for part of one of the notebooks, and Figure 4 presents a montage of all of the generated images in the notebook. The full notebooks are available on Zenodo (https://zenodo.org/records/16033603).

**Figure 3.**
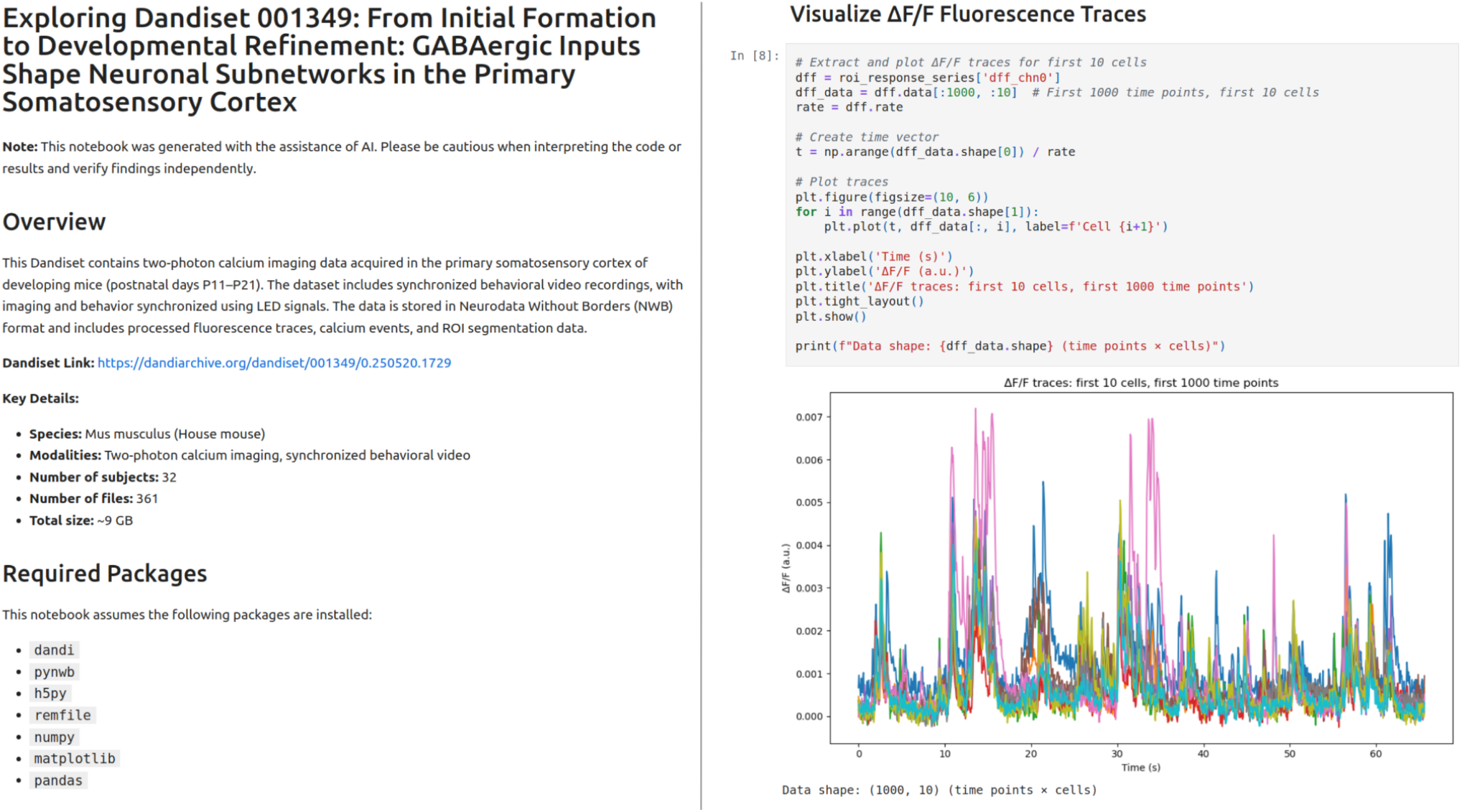
Screenshots from part of the generated notebook for dandiset 001349.

**Figure 4.**
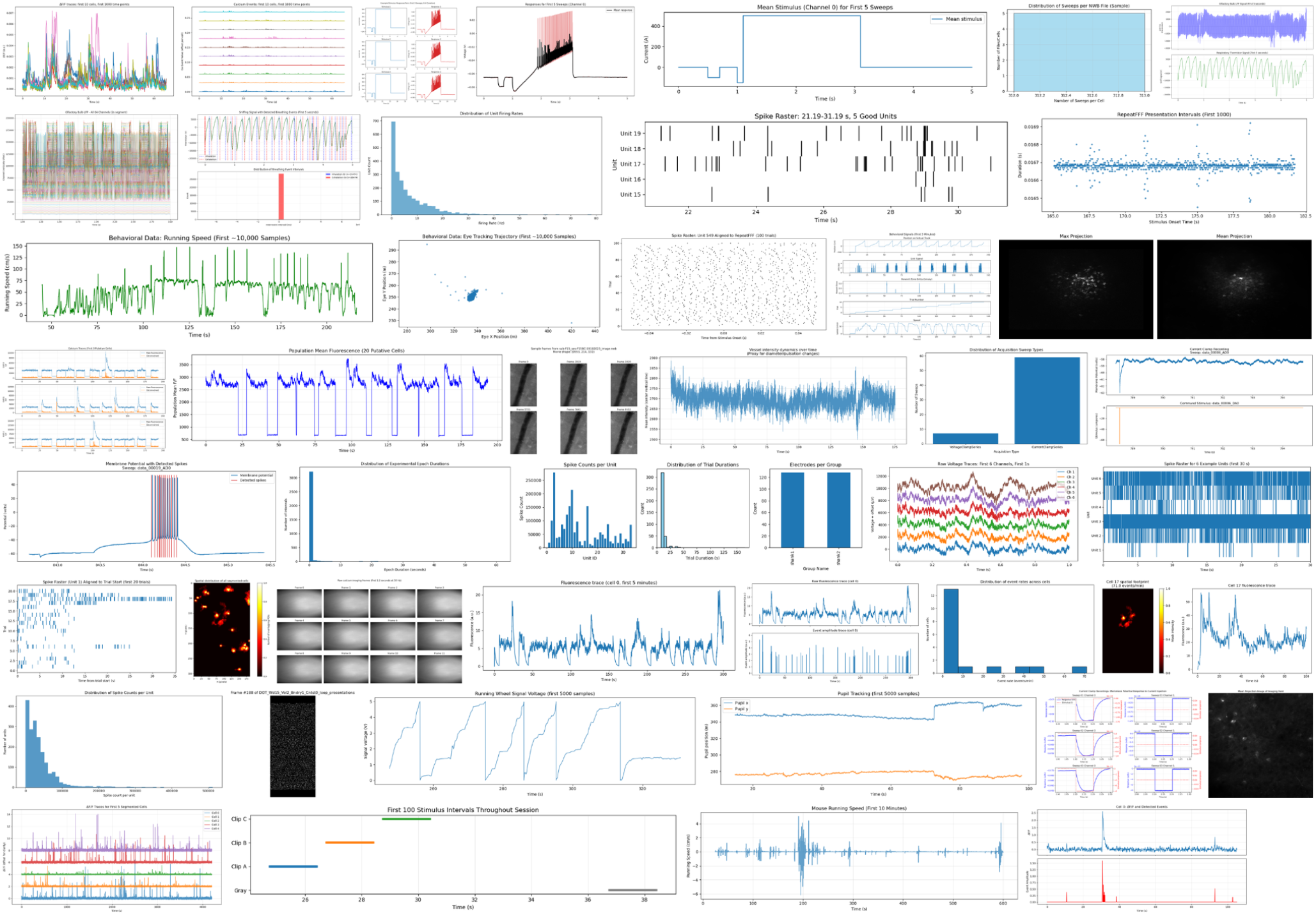
Montage of the 45 images from the 12 AI-generated notebooks.

### Human interventions in the chat phase

User intervention was required in 8 out of 12 dandisets to correct errors or guide the assistant (Table 1). Common issues included misinterpretation of data structures, such as incorrect handling of timestamp fields; ineffective or misleading visualizations, including blank images, sparse raster plots, or misaligned regions of interest (ROIs); and performance problems, such as inefficient data access or code that ran too slowly. For example, for Dandiset 001433, the assistant misinterpreted a dataset of event times, leading to incorrect visualizations. For Dandiset 000617, the assistant repeatedly failed in its attempt to align ROIs with background images, and the user ultimately advised abandoning that approach. For Dandiset 000563, the assistant needed to provide guidance on how to properly load spike trains. Four of the dandisets proceeded without intervention, indicating that the process can operate autonomously in some cases. A key advantage of the chat-based approach is that it provides a clear record that could be used for training evaluation models to correct common errors and enhance prompts to reduce errors.

**Table 1.**
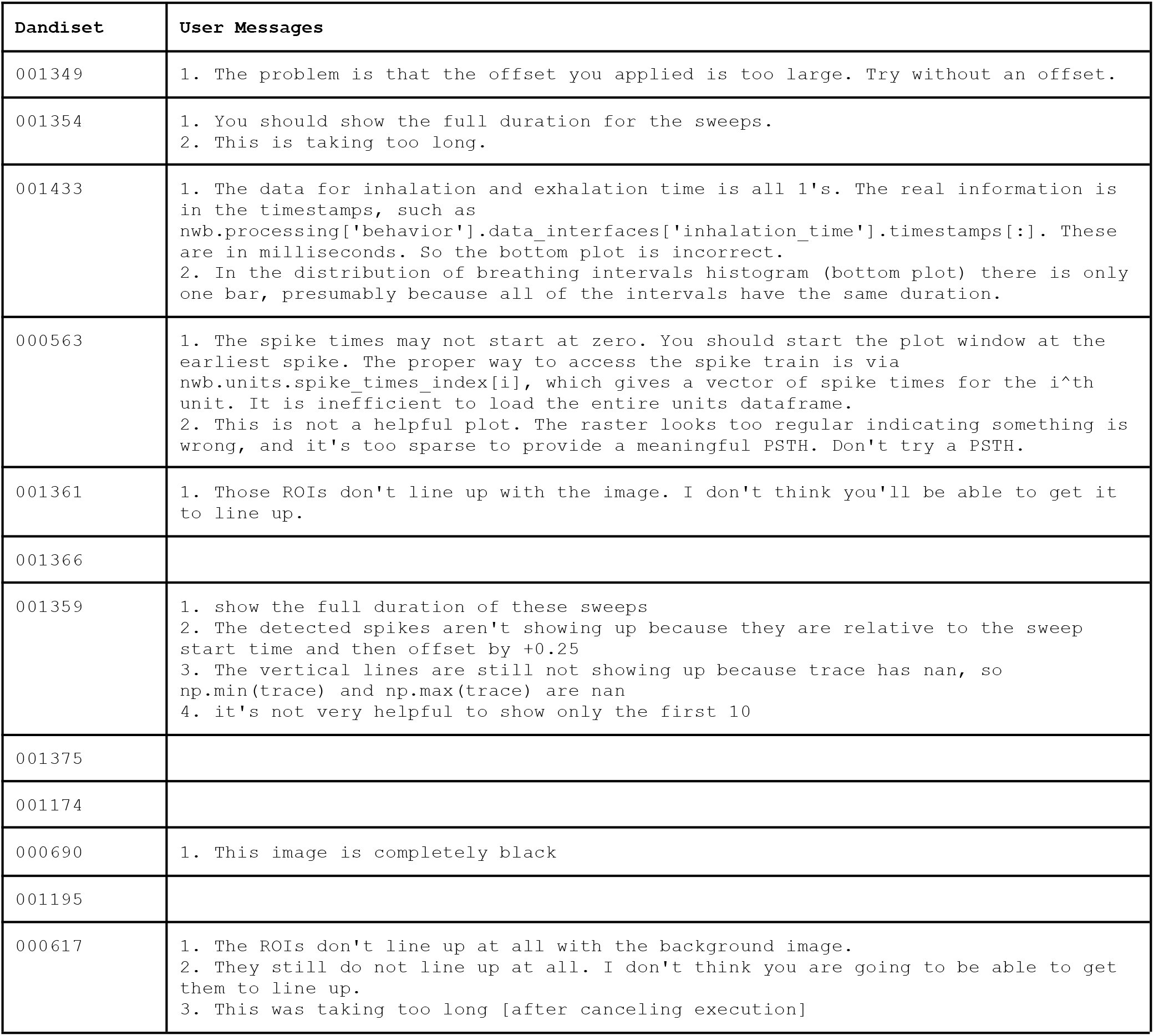
Human interventions in chat conversations.

| Dandiset | User Messages |
| --- | --- |
| 001349 | 1. The problem is that the offset you applied is too large. Try without an offset. |
| 001354 | 1. You should show the full duration for the sweeps.<br>2. This is taking too long. |
| 001433 | 1. The data for inhalation and exhalation time is all 1's. The real information is in the timestamps, such as `nwb.processing['behavior'].data_interfaces['inhalation_time'].timestamps[:]`. These are in milliseconds. So the bottom plot is incorrect.<br>2. In the distribution of breathing intervals histogram (bottom plot) there is only one bar, presumably because all of the intervals have the same duration. |
| 000563 | 1. The spike times may not start at zero. You should start the plot window at the earliest spike. The proper way to access the spike train is via `nwb.units.spike_times_index[i]`, which gives a vector of spike times for the $i^{\text{th}}$ unit. It is inefficient to load the entire units dataframe.<br>2. This is not a helpful plot. The raster looks too regular indicating something is wrong, and it's too sparse to provide a meaningful PSTH. Don't try a PSTH. |
| 001361 | 1. Those ROIs don't line up with the image. I don't think you'll be able to get it to line up. |
| 001366 |  |
| 001359 | 1. show the full duration of these sweeps<br>2. The detected spikes aren't showing up because they are relative to the sweep start time and then offset by +0.25<br>3. The vertical lines are still not showing up because trace has nan, so `np.min(trace)` and `np.max(trace)` are nan<br>4. it's not very helpful to show only the first 10 |
| 001375 |  |
| 001174 |  |
| 000690 | 1. This image is completely black |
| 001195 |  |
| 000617 | 1. The ROIs don't line up at all with the background image.<br>2. They still do not line up at all. I don't think you are going to be able to get them to line up.<br>3. This was taking too long [after canceling execution] |

### Human notebook evaluation

The generated notebooks were independently evaluated by four expert reviewers with backgrounds in data science and neurophysiology and nine reviewers with varying experience with NWB and DANDI. None of these reviewers are authors of this paper. Reviewers completed a structured questionnaire (Table 2) designed to assess the notebooks based on correctness, usefulness, and clarity. The reviewers were given one hour to complete as many notebook reviews as they could in the allotted time and did not discuss the notebooks with anyone until after all their reviews were submitted. Each reviewer reviewed between 5 and 10 notebooks.

**Table 2.** Questions with choices in notebook review survey.

| ID | Question | Options |
| --- | --- | --- |
| understand-dandiset | How well did the notebook help you understand the purpose and content of the Dandiset? | 2: Very well<br>1: Somewhat<br>0: Not at all |
| access-data | After reviewing the notebook, do you feel confident in how to access the different types of data from this Dandiset? | 2: Yes, I know how to access and explore the different types of data<br>1: Somewhat, but I would need more guidance for certain data types<br>0: No, I’m still unsure how to access the data |
| work-with-nwb | Did the notebook help you understand the structure of the NWB file(s) and how to work with them? | 2: Yes, I understand the basics and could explore further on my own<br>1: Somewhat, I still have questions<br>0: No, I feel lost or confused |
| visualization-helpful | Did the visualizations in the notebook generally help you understand key aspects of the data? | 2: Yes, they made important points clear<br>1: Somewhat<br>0: No, the visualizations were generally confusing or unhelpful |
| visualization-issues | Did any of the visualizations make it harder to understand the data (e.g., due to poor formatting, unclear axes, or misleading displays)? | 0: No, the visualizations were clear and helpful<br>-1: One or more visualizations were unclear or hard to interpret<br>-2: Yes, one or more visualizations were misleading, confusing, or had substantial issues |
| visualization-confidence | Do you feel more confident creating your own visualizations of the data after seeing the examples in the notebook? | 2: Yes, I feel equipped to start exploring on my own<br>1: Somewhat, but I’d still need guidance<br>0: No, the notebook didn’t provide helpful examples |
| visualization-structure | How well did the visualizations show the structure or complexity of the data? | 2: Very well - they revealed important patterns or relationships<br>1: Somewhat - they hinted at structure, but lacked depth<br>0: Poorly - they didn’t help me understand the data structure |
| unsupported-interpretations | Were there any interpretations or conclusions in the notebook | 0: No, everything felt reasonable and well-supported |
|  | that felt unclear or not well supported by the data shown? | -1: A few things seemed speculative or unclear<br>-2: Yes, there were several conclusions that didn't seem supported |
| redundant-examples | Did any of the plots or examples feel unnecessarily repetitive or redundant? | 0: No, each plot/example helped add something new<br>-1: A few examples felt a bit repetitive<br>-2: Yes, too many examples seemed to show the same thing |
| exploration-guidance | Did the notebook help you understand what kinds of questions or analyses you could do next with this Dandiset? | 2: Yes, it gave clear ideas for further exploration<br>1: Somewhat, I got a few ideas<br>0: No, it didn't help me think about what to do next |
| neurosift-exploration | Briefly explore the provided Neurosift pages associated with this Dandiset. Were any data types in the explored NWB file not shown or discussed in the notebook? | 0: No, it comprehensively represents the data<br>-1: Yes, it omitted one or a few important data types or features<br>-2: Yes, it omitted several important data types or features |
| neurosift-accuracy | Briefly explore the provided Neurosift pages associated with this Dandiset. Did the notebook inaccurately represent any data? | 0: No, it accurately represents the data as far I can tell<br>-1: Yes, one or a few descriptions or visualizations were incorrect<br>-2: Yes, several descriptions or visualizations were incorrect |
| overall-helpfulness | Overall, how helpful was this notebook for getting started with this Dandiset? | 2: Very helpful<br>1: Moderately helpful<br>0: Not helpful |
| neuroscience-expertise | Your background in neuroscience: | 2: Expert<br>1: Intermediate<br>0: Novice |
| python-expertise | Your experience with data analysis in Python: | 2: Expert<br>1: Intermediate<br>0: Novice |
| nwb-dandi-expertise | Your experience with NWB/DANDI: | 2: Expert<br>1: Intermediate<br>0: Novice |
| prior-dandiset-familiarity | How familiar were you with this Dandiset before reviewing the notebook? | 2: Very familiar<br>1: Somewhat familiar<br>0: Not familiar |
| review-confidence | How confident do you feel in your ability to review this notebook? | 2: Very confident<br>1: Somewhat confident<br>0: Not confident |

Reviewers found the notebooks to be at least moderately effective across most criteria (Figure 5). Most ratings were either “somewhat helpful” or “very helpful,” with very few scores at the lowest end of the scale. Reviewers generally reported success in accessing data, working with NWB files, and interpreting visualizations. There were very few reports of redundancy with the example explorations, and the representation was largely accurate with what was observed independently using Neurosift.

**Figure 5.**
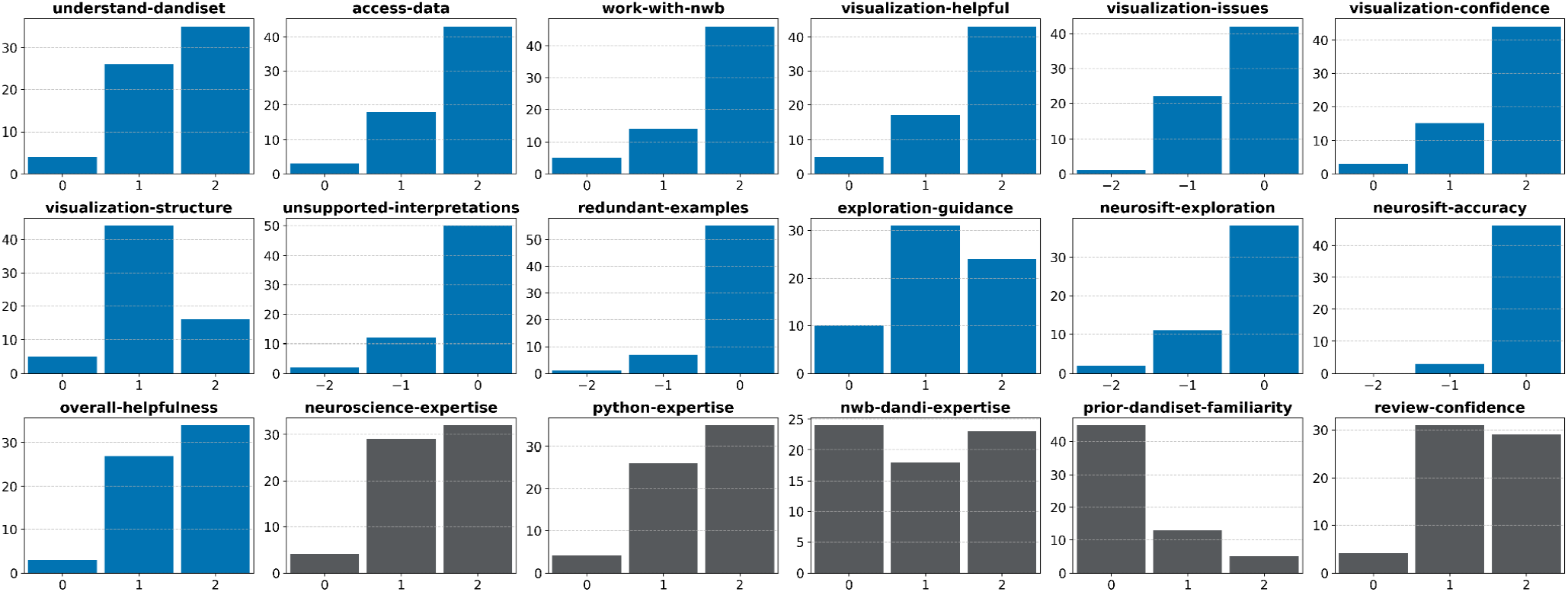
Summary of reviews for AI-generated notebooks. Each subplot shows the distribution of responses to a survey question evaluating aspects such as usefulness, accuracy, visualization quality, and reviewer expertise. The final 5 questions relate to the reviewer’s experience level. See Table 2 for details on the questions and the response options.

Figure 6 shows the breakdown of “Overall Helpfulness” ratings by Dandiset ID and by reviewer experience with NWB and DANDI. Most notebooks received a mix of “moderately helpful” and “very helpful” ratings, with only one dandiset (000690) receiving any “not helpful” responses. Notably, reviewers across different levels of experience with NWB and DANDI gave similar ratings, suggesting that the notebooks are broadly accessible and useful to a wide range of researchers.

**Figure 6.**
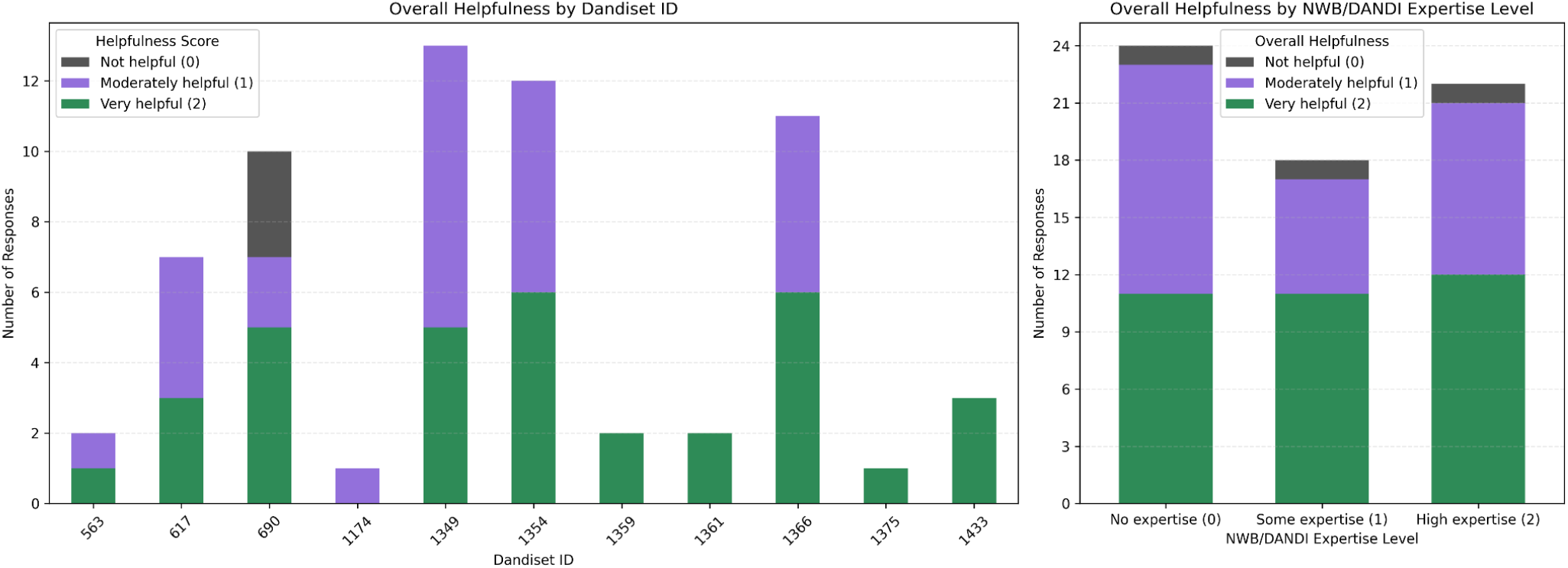
Left: Distribution of “Overall Helpfulness” ratings for each AI-generated notebook. Bars show the number of reviewers who rated each notebook as not helpful (0), moderately helpful (1), or very helpful (2). Note that one notebook was not reviewed by any of the reviewers. Right: Distribution of “Overall Helpfulness” ratings by reported experience with NWB/DANDI.

### LLM cost

We recorded API costs for both the exploration and generation phases of notebook generation across all 12 dandisets. Chat sessions using GPT-4.1 ranged from $0.37 to $1.84 (median $1.03), while notebook generation with Claude Sonnet 4 ranged from $0.09 to $0.46 (median $0.17). The cost for one of the notebooks was not available due to technical issues. The average total cost per notebook was $1.15.

## Discussion

### Reducing barriers to data reuse

This work introduces an AI-powered system that lowers barriers to reusing neurophysiology data from the DANDI Archive by combining LLM-driven automation with lightweight human oversight. The system automates early exploration of NWB datasets and generates ready-to-run, dataset-specific notebooks that help users get started with access, visualization, and analysis. These notebooks are designed to work both locally and in cloud environments such as Google Colab or DANDI Hub. By providing tailored, interactive support, the system enables faster onboarding for both new and experienced users, without requiring detailed knowledge of the NWB data standard or software.

### Enhancing incentives for data sharing

By lowering the technical and conceptual barriers to using open neurophysiology datasets, these tools have the potential to foster new forms of collaboration between data generators and data analysts. When reuse becomes easier and more visible, experimental datasets are more likely to serve as the foundation for secondary analyses and derivative publications that include or credit the data generators. This, in turn, creates a stronger incentive for researchers to adopt standardized formats like NWB and deposit their data in public archives. Future extensions of this work could further support this ecosystem by integrating notebook generation into the data sharing workflow itself, enabling contributors to review and refine AI-generated notebooks before they are published alongside the dataset.

### Balancing automation and human oversight

While the system is largely autonomous, minimal human oversight during the exploration phase was essential for maintaining quality. Interventions were limited to clear issues such as broken visualizations, misread data structures, or flawed analyses rather than guiding the process throughout. These light corrections helped avoid misleading outputs, especially when dataset-specific quirks (e.g., missing units, misaligned ROIs) required expert judgment. Oversight was generally efficient and unobtrusive, but scaling to the full set of 400+ dandisets may require new strategies. Whether future models can fully replace this role remains an open question.

### LLM model selection

We qualitatively assessed several LLMs for the two stages of our pipeline and selected different models for each. OpenAI’s GPT-4.1 was selected for the chat phase, reliably calling tools at the right times, avoiding overinterpretation, and keeping analysis appropriately introductory. For notebook generation, Claude Sonnet 4 was selected, demonstrating strong scientific reasoning and producing creative, well-structured notebooks with high-quality visualizations when constrained to analyses completed during exploration. Claude was less suitable for the exploratory chat phase, where it tended to overanalyze data, generate overly complex plots, and draw unsupported conclusions. Other models, including DeepSeek R1 and Google Gemini 2.0, were excluded due to weaker task adherence and inconsistent tool use. We did not test OpenAI’s most advanced models due to cost. Given the rapid pace of progress in the LLM ecosystem, our choices reflect a pragmatic snapshot of what was available and affordable at the time.

### Generalization to other domains

While this work focuses on neurophysiology and the DANDI Archive, the underlying approach is broadly applicable to other scientific domains. There is nothing inherently unique about neurophysiology that makes this system viable; rather, it relies on two key prerequisites: (1) a library of openly accessible, structured datasets, and (2) standardized tools or APIs for efficient programmatic access. Fields such as genomics, earth sciences, materials science, and astronomy already meet these criteria to varying degrees. The system is particularly well-suited for exploratory analysis tasks that can be executed quickly, as the computational cost of notebook generation scales with execution time. We believe this framework can serve as a general model for accelerating data reuse and lowering entry barriers across a wide range of scientific disciplines.

### Risks and limitations

While LLMs can accelerate scientific analysis, they also pose risks. These include hallucinations (confident but incorrect statements or code), misleading visualizations, overinterpretation of results without proper statistical support, and automation bias where users may trust AI outputs without verification. To reduce these risks, we rely on various safeguards when generating notebooks. A key element is the initial exploration phase, which allows the AI system to learn about the dataset’s structure and content prior to notebook generation. The agent catches and fixes code that produces an error. An expert user reviews code to prevent scientifically incorrect analyses and help ensure trust and reliability of results. This phase not only improves the accuracy of the final notebook but also allows for the detection and correction of metadata inconsistencies or unsupported file types in the source dataset, providing valuable feedback for curation to enhance data quality. Additionally, all notebooks include a clear disclaimer noting they were AI-generated and should be critically reviewed before use.

These notebooks were designed to serve as instructional guides, not to produce novel scientific insights. This narrower objective makes the task considerably more manageable for the AI system. Rather than interpreting raw neurophysiology data to uncover new findings, the agent assembles familiar scientific context and standard analysis code, much of which likely reflects patterns in its training data. By focusing on explanation and usability rather than discovery, the system can reliably generate useful, dataset-specific tutorials without needing deep scientific inference. Whether such systems could eventually support novel research on public neurophysiology data remains an open question for future research.

The effectiveness of our system is inherently constrained by the current capabilities of LLMs. While these models can produce syntactically correct and often helpful code and commentary, their grasp of scientific nuance is limited and occasionally inconsistent. As a result, some generated notebooks may feature superficial analyses, overlook relevant methodological considerations, or offer interpretations that sound plausible but are not rigorously supported. We expect improvements in LLMs will reduce these concerns in the future.

The notebook generator is optimized for datasets that can be explored within a modest computational budget. Many dandisets contain large or complex data files that exceed what is practical to process automatically. To address this, the agent typically selects a single NWB file for inspection and focuses on representative data arrays and metadata that are easily accessible. While this strategy ensures feasibility, it may miss important experimental conditions or data features not present in the sampled file.

## Conclusion

We have presented an AI-powered system that lowers the barrier to analyzing public neurophysiology datasets from the DANDI Archive. By combining a conversational LLM agent with an automated notebook generation pipeline, the system helps users navigate dataset contents, access NWB files, and generate meaningful visualizations. Human review of notebooks demonstrated that the approach can be effective and helpful. While challenges remain, this work illustrates how LLMs can support more accessible, scalable, and reproducible use of open scientific data. The framework may be generalizable to other domains where standardized formats and programmatic access are available.

## Methods

### Model selection

We used OpenAI’s GPT-4.1 for the exploration phase and Anthropic’s Claude Sonnet 4 was used in the notebook generation phase.

### Prompts

Prompts for the LLM agents were iteratively developed through trial and error, based on observed performance across datasets. Prompts for the exploration phase and the notebook generation are provided in Appendix A. The system prompt for the Dandiset Explorer application was omitted here for brevity and is available on our public GitHub repository.

### Agent framework

All AI agents were implemented using custom logic that utilized API calls to OpenRouter [OpenRouter].

### Code execution

Code execution during both phases of notebook creation was performed primarily on DANDI Hub, with some computation being performed on the author’s laptop. Notebook execution was limited to 600 seconds; if a timeout occurred, the LLM agent was notified and a modified version, presumably with reduced data loading, was generated and executed.

### Dandiset selection

Recently published Dandisets were selected for inclusion in this study as described in the Results section.

### Notebook generation

In the second phase of notebook generation, an LLM is tasked with generating a complete Jupyter notebook based on the chat conversation from the first phase (Appendix A, first prompt). This includes summaries, visual outputs, and code produced during exploration. The LLM is guided by structured instructions (Appendix A, second prompt) to produce a Jupytext-formatted Python script [Wouts], which is more natural for LLMs to generate than notebook files. This script is then converted to a standard .ipynb notebook file. The notebook begins with a heading, a brief overview of the dandiset, and a disclaimer reminding users to critically assess AI-generated content. Subsequent sections walk the user through accessing the dataset via the DANDI Python API, exploring its NWB files using PyNWB, and visualizing example data. The script is executed in a controlled environment, and the agent engages in an execution and error correction loop, similar to that of the exploration phase, until a fully functioning notebook is produced without runtime errors. Because the code is adapted from already debugged exploration code, fewer errors and correction cycles are expected.

### Chat application system architecture

The Dandiset Explorer was implemented as a web-based application using React and TypeScript, with a custom chat interface. The system integrates with LLMs through the OpenRouter API. Code execution is handled through integration with Jupyter kernels, allowing real-time Python code execution in either local Jupyter servers or remote JupyterHub instances. The application implements a custom tool system that enables the LLM agents to programmatically interact with the DANDI API for dataset metadata retrieval, access NWB file structure information through a specialized get_nwbfile_info tool (see below), and execute Python scripts with real-time output capture including text, error messages, and generated visualizations. Chat conversations are persisted in a cloud database.

### Agent access to NWB file information

A key component enabling the system’s functionality is a Python tool called get_nwbfile_info, which is available to the Dandiset Explorer agent. This function streams a specified NWB file from the DANDI Archive and extracts detailed information about its contents. It automatically generates usage instructions for accessing each neurodata object in the file and produces structured textual summaries, including object attributes, descriptions, and dataset shapes. This information allows the agent to reason about the dataset structure and generate appropriate code and visualizations during exploration.

## Data and code availability

The AI-generated Jupyter notebooks analyzed in this study are available on Zenodo: https://zenodo.org/records/16033603

The spreadsheet of notebook reviewer scores with accompanying comments/rationale is also available in the same Zenodo record.

The source code for both the notebook generator and the chat, at the time of writing, is available on GitHub: https://github.com/dandi-ai-notebooks/dandi-ai-notebooks-study https://github.com/dandi-ai-notebooks/dandiset-explorer

## Acknowledgments Author Contributions & Competing Interests

This work was supported by the NIH under awards NIH U24NS120057 and NIMH 1R24MH117295.

All authors contributed to the conception and design of the study, software development, data collection and analysis, and writing and reviewing of the manuscript. All authors read and approved the final version.

Benjamin Dichter is the Founder and CEO of CatalystNeuro, a consulting company that specializes in open science in neurophysiology. The other three authors declare no competing interests.

## Appendix A: LLM Prompts

### Initial prompt in the exploration chat

~~~
I would like you to do research on this dandiset to gather information in preparation for
creating a jupyter notebook that introduces this dandiset and helps researchers explore it
and get started with a reanalysis.
Each of your responses should provide information that will be needed in the notebook or
execute code that produces plots or text output that will appear in the notebook. The content
of this conversation will be used to form the notebook, so it should cover everything that is
needed. After you provide each response, I (the user) will either point out any issues with
your response or I will just say “proceed” in which case you will proceed with the research
and exploration. Continue until you have enough information, then say “My research is
complete”. Be thorough.
Do not offer me choices of what to do next. If there are no issues, I will just say “proceed”
and you will continue with the next step of the research.
If you produce a plot and notice there are errors with it that can be remedied, you should
revise your script and regenerate the plot.
The notebook (which will be created based on the content of this conversation) will be evaluated
based on the following criteria:
[CRITERIA ARE LISTED]
~~~

### Prompt for generating the notebook

~~~
On the basis of the above chat, please create a JupyText notebook that introduces dandiset {{
DANDISET_ID }} using the format described below.
The purpose of the notebook is to help researchers explore the dandiset and get started with
a reanalysis.
Start with an appropriate title for the notebook, such as “Exploring Dandiset {{ DANDISET_ID Z
}}: …” (that should be a markdown heading).
Inform the user that the notebook was generated with the assistance of AI, and that they
should be cautious when interpreting the code or results.
Provide an overview of the Dandiset. Include a link to the Dandiset of the form
https://dandiarchive.org/dandiset/{{ DANDISET_ID }}/{{ DANDISET_VERSION }}.
List the packages that are required to run the notebook. Assume that these are already
installed on the user’s system. Do not include any pip install commands in the notebook.
Show how to use the DANDI Python API to information about the dandiset using code similar to
the following:
client = DandiAPIClient()
dandiset = client.get_dandiset(“001333”, “0.250327.2220”)
metadata = dandiset.get_raw_metadata()
print(f”Dandiset name: {metadata[‘name’]}”)
print(f”Dandiset URL: {metadata[‘url’]}”)
Show how to use the DANDI Python API to explore the .nwb files in the dandiset.
Show how to load and visualize data from the dandiset based on the above chat.
Feel free to organize things differently from how they are in the above chat, but do not make
up new information.
Generate good quality plots (without being redundant).
Load NWB data by streaming the remote file (as done in the above chat rather than downloading
it.
Do not use any functionality of PyNWB or dandi that is not covered in the above chat.
You should stick to material that is covered in the above chat and do not hallucinate.
Be mindful of not loading too much data, considering that it will all be streamed from a
remote source.
Throughout the notebook, include explanatory markdown cells that guide the user through the
process.
The notebook should be well-documented, and follow best practices Include comments in code
cells to explain what each step does.
The Jupytext should use ‘# %% [markdown]’ for markdown cells and ‘# %%’ delimiters for the
code cells.
If any NWB files have units objects, you should know the following:
units.spike_times_index[i] provides the vector of spike times for the i^th unit. It is
actually not an index. Do not use units.spike_times.
Do not render or display the NWB object obtained from NWBHDF5IO directly in the notebook as
the output could be very large.
Use concise scientific language.
Prioritize correctness and accuracy over verbosity and complexity.
STRICT RULE: Do not include new analyses that are not covered in the chat conversation.
STRICT RULE: You should learn from any feedback in the conversation to avoid pitfalls in
plots. For example, if the user suggests that a plot is misleading, invalid, or unhelpful,
you should either not include that plot in the notebook or provide a corrected version of the
plot that addresses the user’s concerns. Only provide a corrected version if you are
confident that you can fix the issue.
Do not be verbose in your summary or wrap up of the notebook, although you may briefly
suggest some general ideas for future exploration.
There is no need to close files or clean up resources at the end of the notebook.
Your notebook will be evaluated based on the following criteria:
[CRITERIA ARE LISTED]
Your output must be in the format:
<notebook>
The text of the JupyText notebook should appear here.
</notebook>
No other text should be included in the response.
~~~

